# Modelling the influence of the hippocampal memory system on the oculomotor system

**DOI:** 10.1101/303511

**Authors:** Jennifer D. Ryan, Kelly Shen, Arber Kacollja, Heather Tian, John Griffiths, Gleb Bezgin, Anthony R. McIntosh

**Author notes:** equal contribution (Jennifer D. Ryan, Kelly Shen). Corresponding Authors: Jennifer D. Ryan, PhD, Rotman Research Institute, Baycrest, 3560 Bathurst St., Toronto, ON M6A2E1,; Kelly Shen, PhD, Rotman Research Institute, Baycrest, 3560 Bathurst St., Toronto, ON M6A2E1. **Acknowledgments:** The authors wish to thank R. Shayna Rosenbaum for her helpful comments on a previous version of this manuscript. This work was funded by a grant from Natural Sciences and Engineering Research Council of Canada to J.D.R.

## Abstract

Visual exploration is related to activity in the hippocampus (HC) and/or extended medial temporal lobe system (MTL), is influenced by stored memories, and is altered in amnesic cases. An extensive set of polysynaptic connections exists both within and between the HC and oculomotor systems such that investigating how HC responses ultimately influence neural activity in the oculomotor system, and the timing by which such neural modulation could occur is not trivial. We leveraged TheVirtualBrain, a software platform for large-scale network simulations, to model the functional dynamics that govern the interactions between the two systems in the macaque cortex. Evoked responses following the stimulation of the MTL and some, but not all, subfields of the HC resulted in observable responses in oculomotor regions, including the frontal eye fields (FEF), within the time of a gaze fixation. Modeled lesions to some MTL regions slowed the dissipation of HC signal to oculomotor regions, whereas HC lesions generally did not affect the rapid MTL activity propagation to oculomotor regions. These findings provide a framework for investigating how information represented by the HC/MTL may influence the oculomotor system during a fixation and predict how HC lesions may affect visual exploration.

**Author Summary:** No major account of oculomotor (eye movement) guidance considers the influence of the hippocampus (HC) and broader medial temporal lobe (MTL) system, yet it is clear that information is exchanged between the two systems. Prior experience influences current viewing, and cases of amnesia due to compromised HC/MTL function show specific alterations in viewing behaviour. By modeling large-scale network dynamics, we show that stimulation of subregions of the HC, and of the MTL, rapidly results in observable responses in oculomotor control regions, and that HC/MTL lesions alter signal propagation. These findings suggest that information from memory may readily guide visual exploration, and calls for a reconsideration of the neural circuitry involved in oculomotor guidance.

## Introduction

Memory influences ongoing active exploration of the visual environment (Hannula et al., 2010). For instance, more viewing is directed to novel versus previously viewed items (Fagan, 1970; Fantz, 1964), and more viewing is directed to areas that have been altered from a prior viewing (Ryan, Althoff, Whitlow, & Cohen, 2000; Smith, Hopkins, & Squire, 2006). A number of studies have implicated a network of subregions within the hippocampus (HC) and/or broader medial temporal lobe (MTL) responsible for the influence of memory on viewing behavior. Amnesic cases who have severe memory impairments due to compromised function of the HC and/or MTL show changes in their viewing behavior compared to neurologically-intact cases (Chau, Murphy, Rosenbaum, Ryan, & Hoffman, 2011; Hannula, Ryan, Tranel, & Cohen, 2007; Olsen et al., 2015; Ryan et al., 2000; Warren, Duff, Tranel, & Cohen, 2010). Similar findings have been observed in older adults who have suspected HC/MTL compromise (Ryan, Leung, Turk-Browne, & Hasher, 2007), and certain viewing patterns have been shown to track with entorhinal cortex (ERC) volumes (Yeung et al., 2017). Visual exploration predicts HC activity during encoding (Z.-X. Liu, Shen, Olsen, & Ryan, 2017), and, conversely, HC/MTL activity predicts ongoing visual exploration that is indicative of memory retrieval (Hannula & Ranganath, 2009; Ryals, Wang, Polnaszek, & Voss, 2015). The relationship between visual sampling and HC activity is weakened in aging, presumably due to decline in HC structure or function (Z. X. Liu, Shen, Olsen, & Ryan, 2018). Such evidence collectively demonstrates that HC/MTL function is related to oculomotor *behavior.* The indirect implication of these studies is that the HC must influence *neural activity* in the oculomotor system.

Studies in non-human primates have shown that HC/MTL activity is linked to oculomotor behavior. The activity of grid cells in the ERC are tied to eye position (Killian, Jutras, & Buffalo, 2012), while HC/MTL activity is modulated by saccades (Sobotka, Nowicka, & Ringo, 1997) and fixations (Hoffman et al., 2013; Leonard et al., 2015). How HC/MTL activity traverses the brain to influence the oculomotor system has not been shown to date. The oculomotor system is itself a highly recurrent and distributed network (Parr & Friston, 2017) comprised of cortical and subcortical regions responsible for the execution of a saccade (e.g., frontal eye field, FEF; superior colliculus, SC) as well as regions that exert cognitive control over where the eyes should go (e.g., dorsolateral prefrontal cortex, dlPFC; anterior cingulate cortex, ACC; lateral intraparietal area, area LIP) (Bisley & Mirpour, 2019; Johnston & Everling, 2008). Prior work has speculated as to which regions of the brain may be important for bridging the memory and oculomotor systems (e.g., Meister & Buffalo, 2016; Micic, Ehrlichman, & Chen, 2010), but these discussions were limited to regions examined in isolation. There are no known direct connections between hippocampal subfields and the oculomotor system. Yet, by examining whole-cortex connectivity, we have shown that there is an extensive set of polysynaptic pathways spanning extrastriate, posterior parietal, and prefrontal regions that may mediate the exchange of information between the oculomotor and memory systems (Shen, Bezgin, Selvam, McIntosh, & Ryan, 2016). Given the vast anatomical connectivity within and between the memory and oculomotor systems, trying to discern the functional network involved in bridging them is not a trivial problem to tackle. Specifically, the large and complex contribution of recurrent anatomical connections to the functional dynamics of large-scale brain networks must be considered (Spiegler, Hansen, Bernard, McIntosh, & Jirsa, 2016). One crucial question concerning such functional dynamics is whether HC/MTL activity is able to influence the activity related to the preparation of a saccade. In order to impact ongoing visual exploration, HC/MTL activity would likely need to resolve in the oculomotor system within the time of an average duration of a gaze fixation (∼ 250-400 ms) (Henderson, Nuthmann, & Luke, 2013).

To examine the extent to which HC/MTL activity could influence the oculomotor system, we leveraged a computational modeling and neuroinformatics platform, TheVirtualBrain, and simulated the functional dynamics of a whole-cortex directed macaque network when stimulation is applied to HC and MTL nodes of interest. Critically, we examined whether and when evoked activity culminated in responses in key regions within the oculomotor system. Finally, we observed the extent to which the propagation and timing of such activity was altered following lesions to one or more HC/MTL regions in order to understand the neural dynamics that may underly altered visual exploration in cases of HC/MTL dysfunction, such as in amnesia or aging.

## Results

We modelled the influence of HC/MTL activity on the oculomotor system using a connectome-based approach using TheVirtualBrain (see Methods for details). Following Spiegler and colleagues (Spiegler et al., 2016), we assigned a neural mass model to each node and set each to operate near criticality, which is considered to be the point at which information processing capacity is maximal (G. Deco et al., 2014; Ghosh, Rho, McIntosh, Kötter, & Jirsa, 2008). Nodes were then connected together as defined by a weighted and directed macaque structural connectivity matrix and the distance between them defined by a tract lengths matrix. As we were interested in examining signal propagation *from* the memory system *to* the oculomotor system while taking into account the extensive recurrent connectivity between them, we chose to use the macaque connectome because of the available information from tracer data on the directionality of fiber tracts. Without stimulation, this network exhibits no activity. However, with stimulation, activity dissipates throughout the network according to the spatiotemporal constraints imposed by the connectivity weights and distances. Despite having no spontaneous activity, this model has been shown to exhibit the emergent properties of spontaneous activity (Spiegler et al 2016). That is, with stimulation, the model produces a diverse set of resting-state networks that are typically detected from spontaneous activity in empirical studies. Cortical network dynamics were set via additional parameter tuning such that stimulating V1 resulted in biologically plausible timing of evoked responses in downstream visual cortical regions. Finally, we systematically stimulated HC subfields and MTL regions of interest and detected evoked responses across the rest of the network.

### HC Subregion Stimulation

Stimulation of HC subfields and MTL regions of interest evoked widespread activation across the network, similar to previous surface-based model simulations (Spiegler et al., 2016). Figure 1 shows an example of activity dissipation following CA1 stimulation. Evoked responses were first detected in other HC subfields and MTL regions but then spread to prefrontal and extrastriate cortices, and later to posterior parietal cortex. The full list of activation times for each of the 77 nodes can be found in Supplementary Table 2. However, in all subsequent analyses, we present only the results pertaining to our nodes of interest, identified as those along the shortest paths between HC/MTL and oculomotor regions (Shen et al., 2016) or those that have been specifically suggested in the literature to be potentially relevant (Meister & Buffalo, 2016).

**Figure 1.**
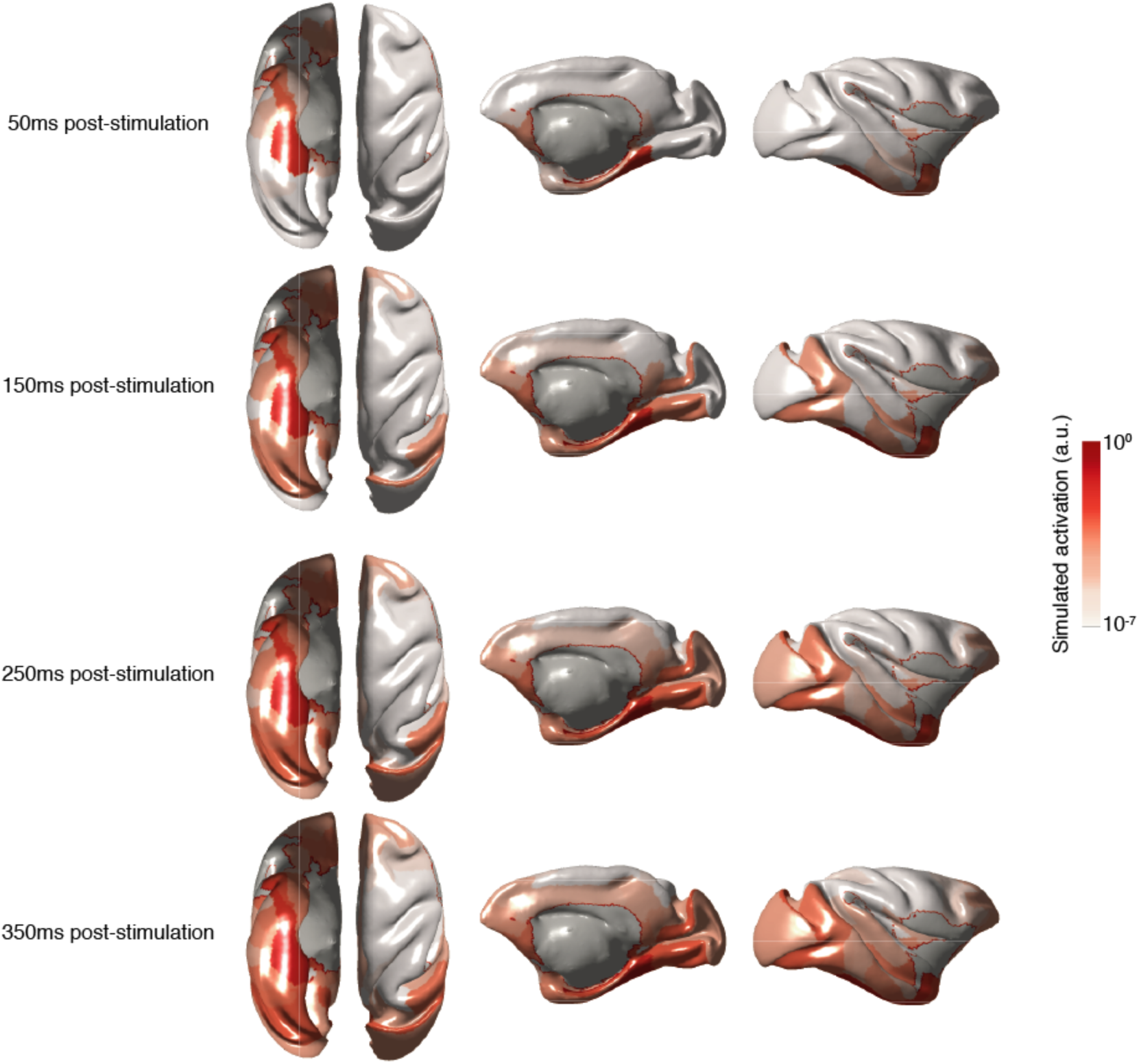
Dissipation of activity over time across the cortex following simulated stimulation of CA1. Average above-threshold simulated activation (arbitrary units) for each node for a 10-ms epoch following each time point is plotted on the macaque cortical surface. Activations were log scaled for the purposes of visualization. From left to right: ventral, dorsal, medial and lateral views of the macaque cortical surface.

Stimulation of HC subregions CA1, subiculum (S), pre-subiculum (PrS), and para-subiculum (PaS) resulted in observable responses in almost all of the cortical nodes of interest, and within regions 46, 24, and FEF, of the oculomotor system (for CA1 example, see Figure 2A). Within our oculomotor regions of interest, activity was first observed in area 46, followed by 24, and FEF, regardless of HC stimulation site. Stimulation of the PrS resulted in the fastest observable responses in these oculomotor areas (under 70 ms; Figure 3A). Stimulation of CA1 resulted in rapid activity that culminated in oculomotor regions in under 220 ms (Figure 3B). Stimulation of either the S or the PaS resolved into area 46 activity by 200 ms, into area 24 by 250 ms, and finally into FEF by 500 ms (Figure 3C-D). Responses in area LIP occurred substantially later than the other oculomotor areas and even later than all other cortical nodes for CA1 and PrS stimulations (> 440 ms; Table 1). No evoked response was detected in area LIP following stimulation of S or PaS. Responses were not observed in the majority of the pre-defined cortical hubs following CA3 stimulation, and activity did not culminate in observable responses in the oculomotor areas (Figure 2B). See Table 1 for activation times for all nodes of interest.

**Figure 2.**
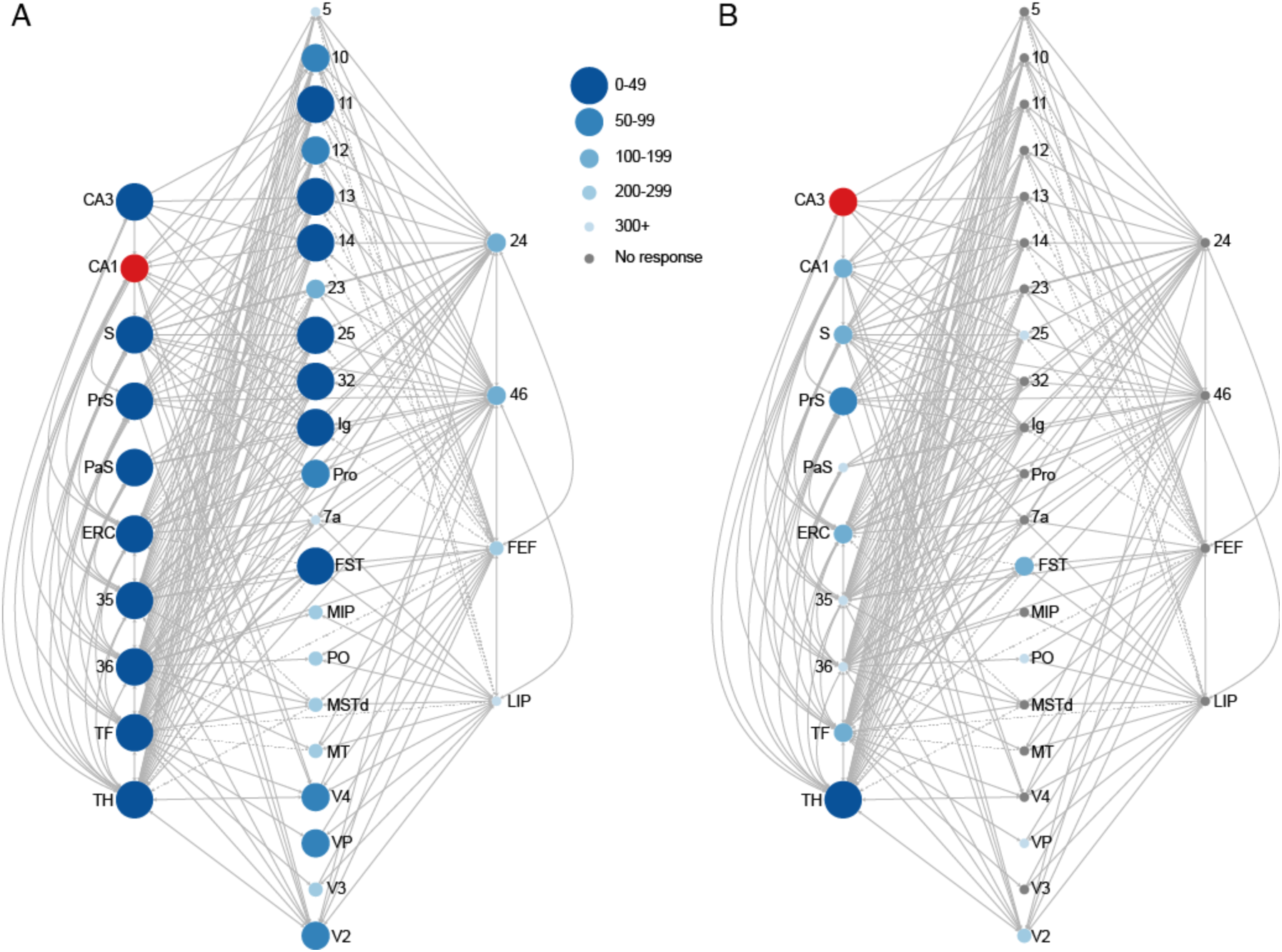
(A) Simulated stimulation of the CA1 (red circle) resulted in observable responses (blue circles) in multiple HC/MTL nodes, intermediary nodes, and in regions governing oculomotor control, including the frontal eye fields (FEF). (B) Simulated stimulation of the CA3 (red circle) resulted in observable responses (blue circles) limited to HC/MTL nodes. Very few responses were observed in cortical areas and none were observed in oculomotor areas. Size and shade of the circles scale with elapsed time prior to an observed response. Grey lines denote direct structural connections between nodes. For visualization purposes, only regions that contribute to the shortest paths *from* HC/MTL *to* oculomotor nodes are shown. Connections between intermediary nodes (middle column) are not shown. Connections that are unidirectional and *away from* oculomotor areas (i.e., to HC/MTL) are indicated by dashed lines.

**Figure 3.**
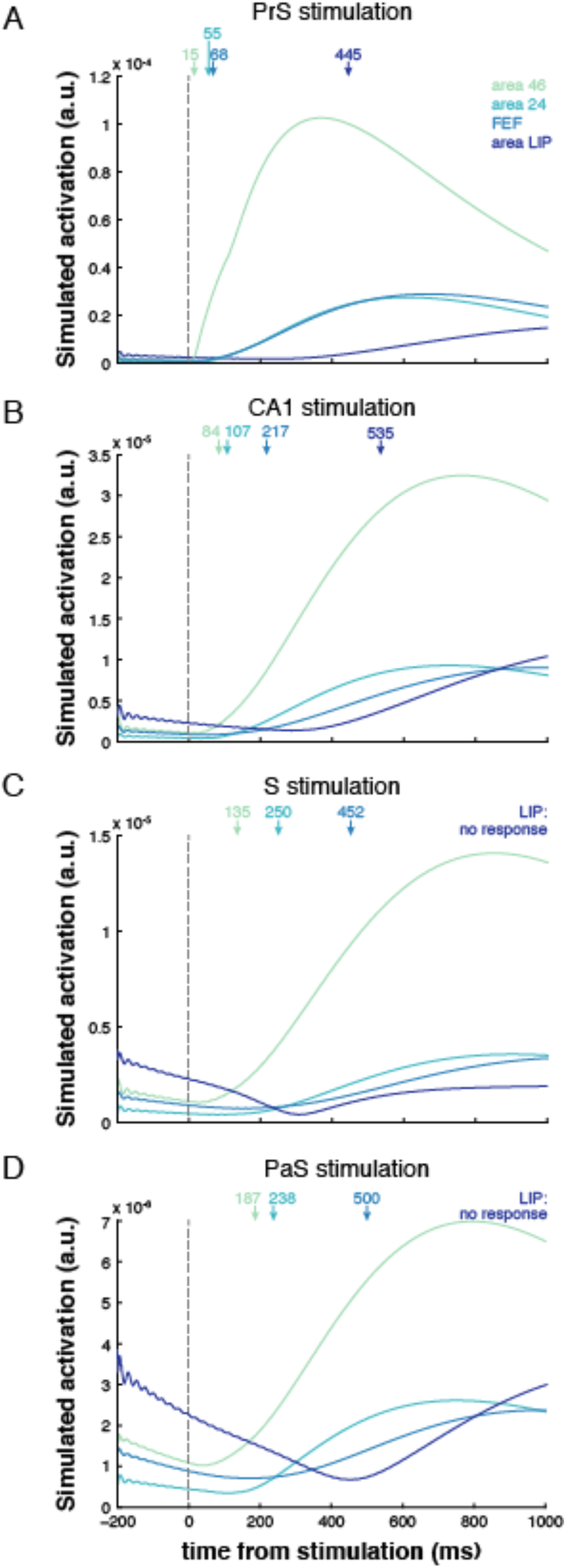
Simulated response profiles (envelope of region time series) of oculomotor areas following stimulation of PrS (A), CA1 (B), S (C) and PaS (D). Activation is given in arbitrary units (a.u.). The onsets of the responses for each oculomotor area indicated by arrows. Area LIP did not exhibit a response that exceeded its baseline threshold following S and PaS stimulation.

**Table 1.**
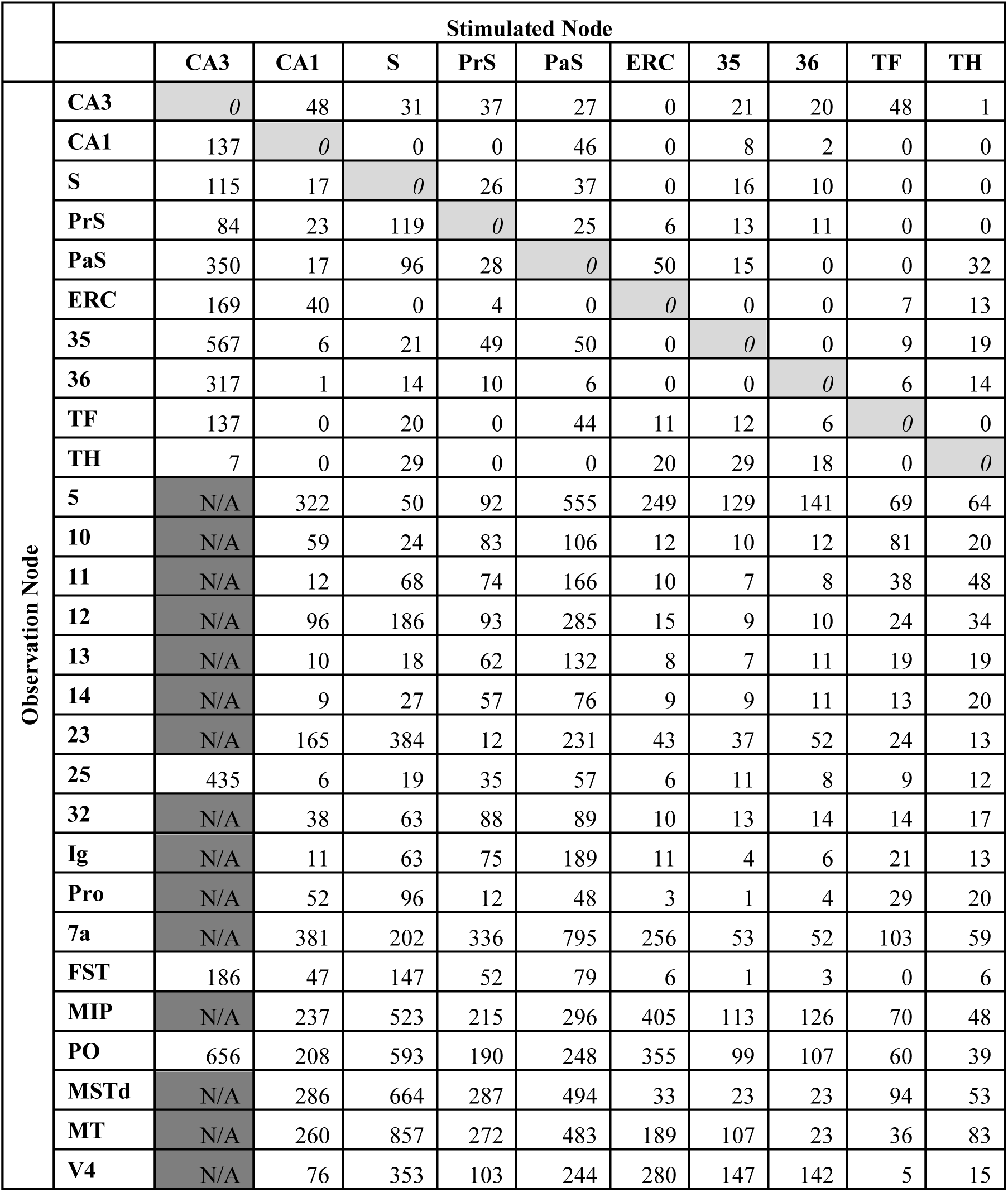
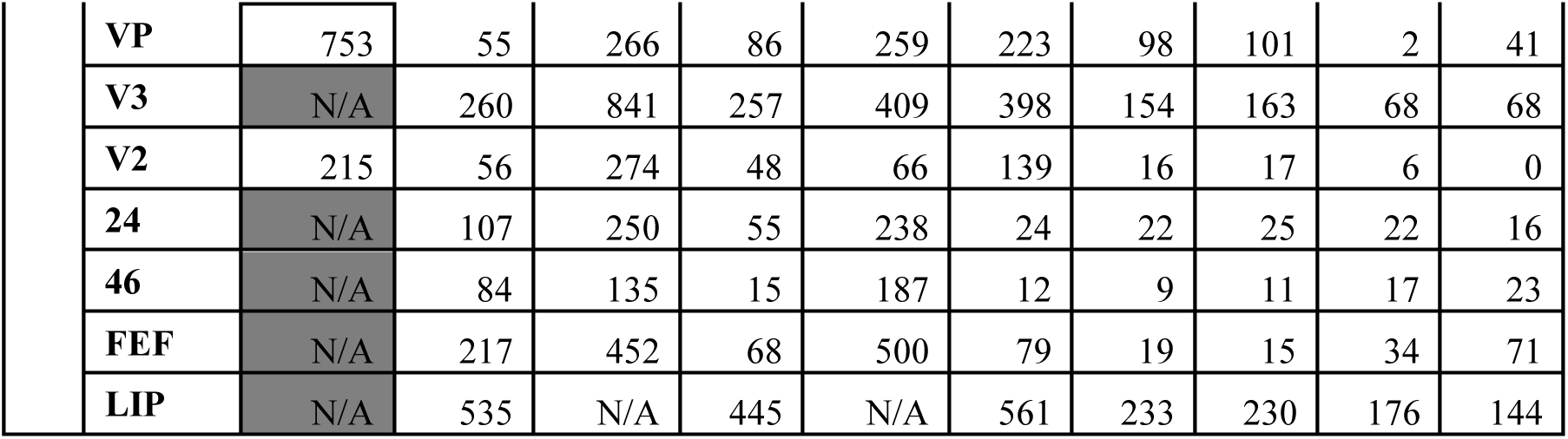
Simulated activation times (ms) following stimulation of hippocampal subfields and medial temporal lobe regions. Only nodes of interest (HC/MTL regions, oculomotor regions, and regions that are involved in the shortest paths between HC/MTL and oculomotor nodes) are shown. S= subiculum, PrS = pre-subiculum, PaS=para-subiculum; ERC = entorhinal cortex; 35/36 = perirhinal cortex; TF/TH = parahippocampal cortex. *0 =* stimulation onset; N/A = no response observed. For a comprehensive set of activation times for all nodes in the network, see Supplementary Table 2.

|  | Stimulated Node |  |  |  |  |  |  |  |  |  |  |
| --- | --- | --- | --- | --- | --- | --- | --- | --- | --- | --- | --- |
|  |  | CA3 | CA1 | S | PrS | PaS | ERC | 35 | 36 | TF | TH |
| Observation Node | CA3 | 0 | 48 | 31 | 37 | 27 | 0 | 21 | 20 | 48 | 1 |
|  | CA1 | 137 | 0 | 0 | 0 | 46 | 0 | 8 | 2 | 0 | 0 |
|  | S | 115 | 17 | 0 | 26 | 37 | 0 | 16 | 10 | 0 | 0 |
|  | PrS | 84 | 23 | 119 | 0 | 25 | 6 | 13 | 11 | 0 | 0 |
|  | PaS | 350 | 17 | 96 | 28 | 0 | 50 | 15 | 0 | 0 | 32 |
|  | ERC | 169 | 40 | 0 | 4 | 0 | 0 | 0 | 0 | 7 | 13 |
|  | 35 | 567 | 6 | 21 | 49 | 50 | 0 | 0 | 0 | 9 | 19 |
|  | 36 | 317 | 1 | 14 | 10 | 6 | 0 | 0 | 0 | 6 | 14 |
|  | TF | 137 | 0 | 20 | 0 | 44 | 11 | 12 | 6 | 0 | 0 |
|  | TH | 7 | 0 | 29 | 0 | 0 | 20 | 29 | 18 | 0 | 0 |
|  | 5 | N/A | 322 | 50 | 92 | 555 | 249 | 129 | 141 | 69 | 64 |
|  | 10 | N/A | 59 | 24 | 83 | 106 | 12 | 10 | 12 | 81 | 20 |
|  | 11 | N/A | 12 | 68 | 74 | 166 | 10 | 7 | 8 | 38 | 48 |
|  | 12 | N/A | 96 | 186 | 93 | 285 | 15 | 9 | 10 | 24 | 34 |
|  | 13 | N/A | 10 | 18 | 62 | 132 | 8 | 7 | 11 | 19 | 19 |
|  | 14 | N/A | 9 | 27 | 57 | 76 | 9 | 9 | 11 | 13 | 20 |
|  | 23 | N/A | 165 | 384 | 12 | 231 | 43 | 37 | 52 | 24 | 13 |
|  | 25 | 435 | 6 | 19 | 35 | 57 | 6 | 11 | 8 | 9 | 12 |
|  | 32 | N/A | 38 | 63 | 88 | 89 | 10 | 13 | 14 | 14 | 17 |
|  | Ig | N/A | 11 | 63 | 75 | 189 | 11 | 4 | 6 | 21 | 13 |
|  | Pro | N/A | 52 | 96 | 12 | 48 | 3 | 1 | 4 | 29 | 20 |
|  | 7a | N/A | 381 | 202 | 336 | 795 | 256 | 53 | 52 | 103 | 59 |
|  | FST | 186 | 47 | 147 | 52 | 79 | 6 | 1 | 3 | 0 | 6 |
|  | MIP | N/A | 237 | 523 | 215 | 296 | 405 | 113 | 126 | 70 | 48 |
|  | PO | 656 | 208 | 593 | 190 | 248 | 355 | 99 | 107 | 60 | 39 |
|  | MSTd | N/A | 286 | 664 | 287 | 494 | 33 | 23 | 23 | 94 | 53 |
|  | MT | N/A | 260 | 857 | 272 | 483 | 189 | 107 | 23 | 36 | 83 |
|  | V4 | N/A | 76 | 353 | 103 | 244 | 280 | 147 | 142 | 5 | 15 |

|  |  |  |  |  |  |  |  |  |  |  |
| --- | --- | --- | --- | --- | --- | --- | --- | --- | --- | --- |
| **VP** | 753 | 55 | 266 | 86 | 259 | 223 | 98 | 101 | 2 | 41 |
| **V3** | N/A | 260 | 841 | 257 | 409 | 398 | 154 | 163 | 68 | 68 |
| **V2** | 215 | 56 | 274 | 48 | 66 | 139 | 16 | 17 | 6 | 0 |
| **24** | N/A | 107 | 250 | 55 | 238 | 24 | 22 | 25 | 22 | 16 |
| **46** | N/A | 84 | 135 | 15 | 187 | 12 | 9 | 11 | 17 | 23 |
| **FEF** | N/A | 217 | 452 | 68 | 500 | 79 | 19 | 15 | 34 | 71 |
| **LIP** | N/A | 535 | N/A | 445 | N/A | 561 | 233 | 230 | 176 | 144 |

### MTL Stimulation

Stimulation of any of the broader regions within the MTL (entorhinal cortex, ERC; perirhinal cortex, 35, 36; parahippocampal cortex, TF, TH) resulted in observable responses within oculomotor areas 46, 24, and FEF well under 100ms, faster than the responses observed from HC subfield stimulation. Of the MTL regions, stimulation of area 35/36 resulted in the earliest responses in areas 46, 24, and FEF (within 25 ms). Although evoked responses in area LIP occurred in under ∼250 ms for all but ERC stimulations, area LIP again exhibited the most delayed response across all nodes of interest following MTL stimulations. See Table 1 for activation times for all nodes of interest.

### Cortical Responses

HC and MTL region stimulation (except for CA3) resulted in signal propagation across all of our pre-identified cortical regions of interest. When CA3 was stimulated, cortical responses were only observed in areas V2 and 25; no other signal was observed. Notably, responses in areas 5 and 7a were generally observed *following* activity from oculomotor regions, including FEF, suggestive of a possible feedback response. The exception is S stimulation, in which responses in area 5 preceded responses in oculomotor regions by ∼100 ms. Responses in V4 also followed oculomotor responses; except in cases of CA1, TF, and TH stimulation. Likewise, responses in area 23 followed oculomotor responses, except in cases of PrS and TH stimulation. See Table 1 for activation times for all nodes of interest.

### Lesion Models: HC subregions

Some models of HC and MTL lesions showed an appreciable effect on activation times while others did not. Only the results for lesions that affected any activation time by at least ±10 ms are shown. Lesion of CA3 changed neither the pattern nor the timing of observable responses following stimulation of each of the other HC/MTL regions (data not shown). Lesion of CA1 resulted in a lack of signal to V2, V4, area 23, and slowing of signal from the subicular complex to various regions, including oculomotor regions FEF and area 24 (Figure 4A; Supplementary Table 3). Lesion of CA1 also led to small increases in the speed of signal following CA3 stimulation to the subicular complex, and from MTL regions to TF/TH, and to other regions within the subicular complex (all less than 10 ms).

**Figure 4.**
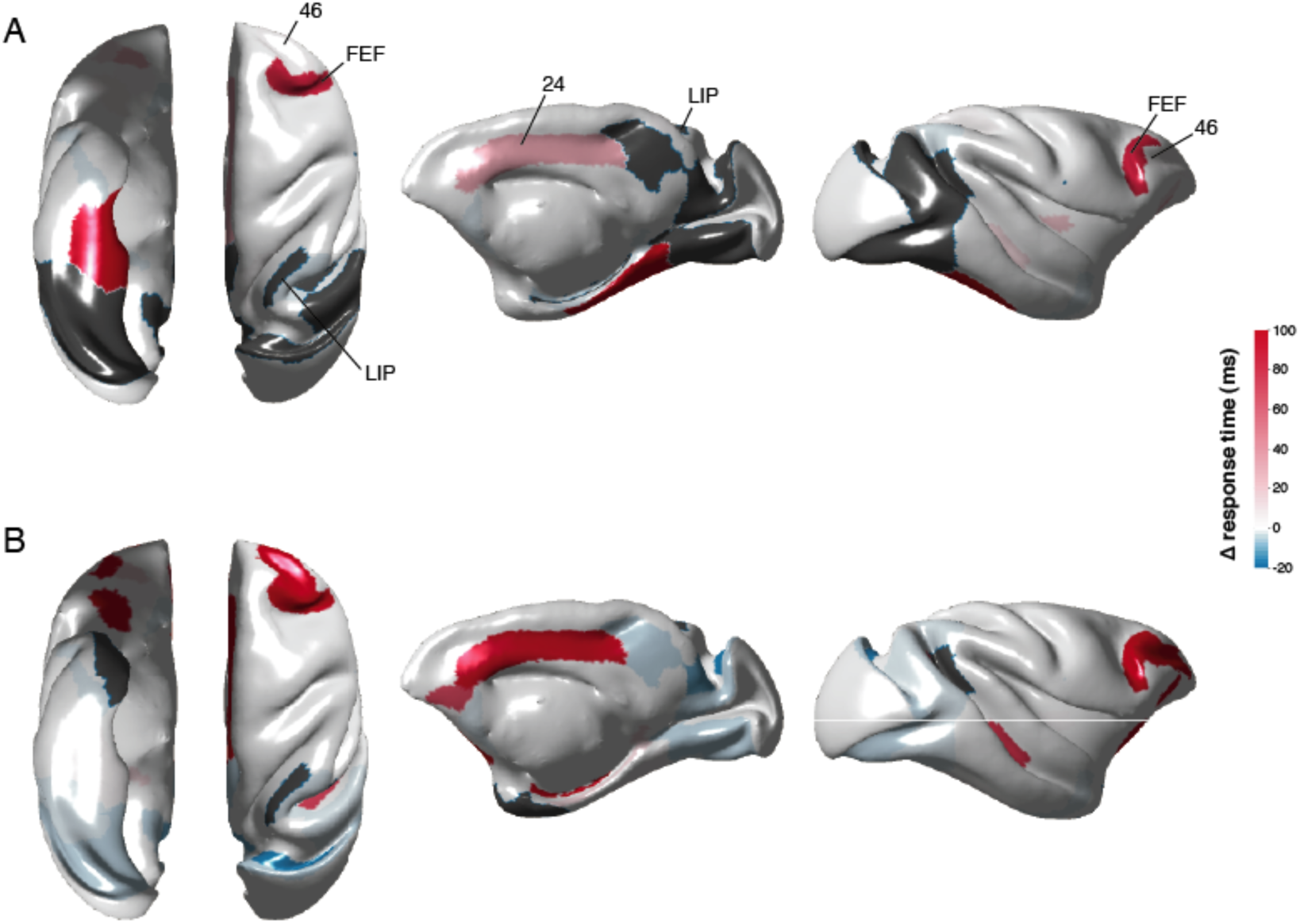
Changes in simulated activation times following HC lesions. Subicular stimulation following CA1 (A) and ERC (B) lesions. Only nodes of interest are presented on the brain surface plots. Activation time differences were computed by subtracting the pre-lesion activation times from the post-lesion ones. Absence of response following a lesion indicated in grey. From left to right: ventral, dorsal, medial and lateral views of the macaque cortical surface.

Lesions to either the S or PaS produced little change to either the pattern or timing of responses following stimulation of the other HC/MTL regions (data not shown). Lesions to the PrS produced moderate changes (<20 ms) in timing: there was some slowing of activity propagation from PaS to some cortical regions, including oculomotor regions, and some speeding of signal propagation within the HC subfields and to TF/TH (Supplementary Table 4).

A combined lesion to all HC subfields (CA3/CA1/S/PaS/PrS) did not considerably change the pattern of signal propagation from the MTL cortices to oculomotor regions. In cases where speeding/slowing was observed, the timing differences were less than 15 ms, and mostly less than 10ms. Signal from the MTL cortices still culminated within the oculomotor regions well under 50 ms (except for ERC -> FEF at 79 ms, and LIP whose responses remained >140 ms) (Supplementary Table 5).

### Lesion Models: MTL regions

Lesion of the ERC resulted in considerable slowing of observable signal in areas 24, 46 and FEF (30-340ms) following S (Figure 3B) or PaS stimulation (Supplementary Table 6). TF and/or TH lesions resulted in slowing (10-400ms) of signal following CA1, S, and PaS stimulation to one or more of areas 24, 46 and FEF, and a lack of response in FEF following PaS stimulation (only the combined TF/TH is shown; Supplementary Table 7). <u>A</u>rea 35 and/or 36 lesions also resulted in slowing (10-90ms) of signal following CA1, S, and PaS stimulation to one or more of areas 24, 46 and FEF, although not as severe as the slowing observed following TF/TH lesions (only the combined 35/36 lesion is shown; Supplementary Table 8).

### Other Cortical Lesions

In our original stimulations, signals in regions 5, 7a, 23, and V4 were predominantly observed following observable responses in oculomotor areas 24, 46 and FEF, suggesting these cortical areas are receiving feedback signals rather than primarily serving as hubs to transfer signal from the HC/MTL to the oculomotor regions. To explore this in more depth, we simulated a combined lesion of 5/7a/23/V4 and examined signal propagation. Following this combined cortical lesion, stimulation of each of the HC/MTL regions (except for CA3) continued to result in observable signal in areas 24, 46 and FEF. Interestingly, while area LIP exhibited the slowest overall responses in the intact simulations, the combined cortical lesion led to substantial speeding of signal to LIP from HC (200-270 ms faster) and MTL (80-300 ms faster) regions owing to speeded responses in intermediary visual cortical and parietal areas (Supplementary Table 9).

## Discussion

A preponderance of evidence has demonstrated a correlation between HC/MTL neural activity and oculomotor behavior (Hannula et al., 2010; Killian, Potter, & Buffalo, 2015; Z.-X. Liu et al., 2017), but research had not shown whether HC/MTL activity can reach the oculomotor system in time to influence the preparation of a saccade. The HC is well connected anatomically to the oculomotor system through a set of polysynaptic pathways that span MTL, frontal, parietal, and visual cortices (Shen et al., 2016) but the existence of anatomical connections does not provide conclusive evidence of the functional relevance of specific pathways. By considering the functional dynamics and recurrent interactions of the large-scale network involved in the HC/MTL guidance of eye movements, we show that propagation of evoked HC/MTL neural activity results in neural activity observable in areas 24 (ACC), 46 (dlPFC) and FEF, which are important for the cognitive and motoric control of eye movements, respectively (Johnston & Everling, 2008). Critically, the culmination of neural signal in these oculomotor regions occurred within the time of a typical gaze fixation (∼250-400 ms; Buswell, 1935; Henderson et al., 2013): within 200 ms following HC subfield stimulation (except for CA3), and within 100 ms following stimulation of each MTL region. Our findings suggest that the underlying neural dynamics of the memory and oculomotor systems allow for representations mediated by the HC/MTL to guide visual exploration – what is foveated and when – on a moment-to-moment basis.

The lack of responses in the FEF following CA3 stimulation is not surprising, given that there are no known direct connections, and fewer polysynaptic pathways, between the CA3 and the oculomotor regions investigated here (Shen et al., 2016). These functional and anatomical differences align well with the purported representational functions of CA3 versus CA1. Foveated information may be bound into detailed memory representations via the auto-associative network of the CA3 (*pattern separation*; Norman & O’Reilly, 2003; Yassa & Stark, 2011), whereas CA1 would enable the comparison of stored information to the external visual world (*pattern completion*; Rolls, 2013; Yassa & Stark, 2011).

Stimulation of the subiculum and parasubiculum resulted in relatively slower responses observed in each of the oculomotor regions, whereas stimulation of presubiculum resulted in rapid responses observed in the oculomotor regions. The subiculum and parasubiculum may largely provide information that supports the grid cell mapping of the ERC (Boccara et al., 2010; Peyrache, Schieferstein, & Buzsáki, 2017; Tang et al., 2016). These regions may then function as a ‘pointer’ by providing online information of an individual’s location in space (Tang et al., 2016). This slowly changing spatial layout may not then require a rapid influence on the oculomotor system, but instead, may allow for the presubiculum, which has cells that are responsive to head direction (Robertson, Rolls, Georges-François, & Panzeri, 1999) to precisely locate and foveate visual objects. These functional distinctions are speculative, and remain to be tested.

Stimulation of each of the MTL cortices resulted in observable responses in each of areas 24, 46 and FEF that were faster than any of the responses observed following HC subregion stimulation. The MTL cortices are intermediary nodes that may permit the relatively rapid transfer of information from HC to the oculomotor system. The unique representational content supported by each region may influence ongoing visual exploration in a top-down manner. The PRC provides lasting information regarding the features of objects (Erez, Cusack, Kendall, & Barense, 2016; Graham, Barense, & Lee, 2010), the PHC provides information regarding the broader spatial environment (Alvarado & Bachevalier, 2005; Eichenbaum, Yonelinas, & Ranganath, 2007; Sato & Nakamura, 2006), and the ERC may provide information regarding the relative spatial arrangements of features within (Yeung et al., 2017), and among objects within the environment (Buckmaster, 2004; Yeung et al., 2019). Signal from the MTL may be used to accurately, and rapidly, prioritize gaze fixations to areas of interest.

HC subfield lesions only minimally altered the timing of activity from MTL to oculomotor regions; the relatively rapid propagation of signal from MTL to FEF (< 100ms) was preserved. Lesions to MTL regions resulted in slowing of signal from some HC subfields to oculomotor regions. This pattern of results suggests that different patterns of visual exploration (i.e., rate, area) may occur in cases of HC/MTL damage depending on the location of the lesion. Lesions restricted to HC subfields may result in an increase in the rate of gaze fixations due to the intact, rapid responses from the MTL to oculomotor regions. This is consistent with prior work in which a developmental amnesic case with HC subfield volume reductions showed an increase in gaze fixations compared to control participants (Olsen et al., 2015). Similarly, older adults, who had functional changes in the HC, showed increases in visual exploration (Z.-X. Liu et al., 2017). ERC and PHC lesions may slow the use of information regarding the broader, ongoing spatial environment; this could result in the need to continually revisit regions to re-establish the relations within and among objects, and with their broader environment, and thus an increased area of visual exploration and/or increase between-object gaze transitions would be observed. Such behavior has been shown by older adults, which may be related to structural and/or functional changes in the ERC (Chan, Kamino, Binns, & Ryan, 2011; Yeung et al., 2017, 2019).

A future question for investigation is how distinct types of representations from the HC/MTL are integrated and prioritized to influence visual exploration, including saccade timing and the ordering of gaze fixations to distinct targets. Alternatively, the functionally distinct representations of the HC/MTL may not actually be integrated within the oculomotor system; rather, each may guide visual exploration at different moments, as time unfolds, as new information in the visual world is sampled, and as task demands are enacted and ultimately met. In either case, multiple memory ‘signals’ are evident within patterns of gaze fixations, including memory for single stimuli (Althoff & Cohen, 1999; Smith & Squire, 2017), memory for the relative spatial (and non-spatial) relations within (Yeung et al., 2017), and among objects (Hannula et al., 2007; Ryan et al., 2000; Smith et al., 2006). Dissociations in these memory signals can be observed within single gaze patterns in neuropsychological cases (Ryan & Cohen, 2003).

Memory may influence visual exploration through multiple routes. Responses emanating from the HC/MTL that ultimately resulted in observable responses in the ACC, dlPFC, and FEF traversed through multiple frontal, visual, and parietal nodes. Yet, despite previous speculation for the involvement of area LIP in bridging the memory and oculomotor systems (Meister & Buffalo, 2016), we found no evidence that the functional dynamics operating within the constraints of the macaque connectome could support this notion. Additionally, responses were observed *following* responses in the oculomotor regions in regions 5, 7a, 23 (posterior cingulate), and V4, suggesting that they may receive feedback from oculomotor regions, rather than serving as hubs that relay information between the HC/MTL and oculomotor systems (Meister & Buffalo, 2016). Area 5 has been implicated in mapping visual and body-centered frames of reference to support visually-guided reaching (Seelke et al., 2012). Cells in area 7a are responsive to eye position and saccades (Bremmer, Distler, & Hoffmann, 1997). The posterior cingulate is part of the default mode network (Buckner, Andrews-Hanna, & Schacter, 2008; Vincent, Kahn, Van Essen, & Buckner, 2010) that is active during internally directed cognitions. Neurons in V4 of the macaque are known to integrate visual and oculomotor information, and show remapping of space towards that of a saccade target, thereby bridging pre- and post-saccade spatial representations (Neupane, Guitton, & Pack, 2016). Information from the HC/MTL may guide gaze selection and execution, and the resulting spatial selections are continually updated throughout cortex to promote ongoing exploration, and feed back into memory.

Here, we have discussed the interactions between the oculomotor and HC/MTL systems within the broader context of ‘memory’, due to the wealth of data showing the influence of memory on visual exploration, and the changes to visual exploration that occur due to dysfunctions of memory, such as in amnesia. However, it is important to note that the lasting representations that are mediated by the HC and MTL may be used in service of cognitive operations beyond memory, to include perception, attention, problem-solving, etc. (Cohen, 2015; Graham et al., 2010). Open questions remain regarding when, during formation, the representations that are mediated by the HC/MTL may be stable enough to influence active vision (Chau et al., 2011; Ryan, J.D., Cohen, 2004; Wynn, Ryan, & Moscovitch, 2019). Likewise, while the results from the model here suggest that information emanating from the HC/MTL may influence activity in oculomotor regions within the time of a gaze fixation, there may be differences in the prioritization and timing by which, and even whether, the distinct HC/MTL representations influence ongoing visual exploration depending on the task that is presented to the viewer. Additional empirical evidence is needed to explore these issues.

It should be noted that future work also remains to examine signal propagation across subcortical-cortical pathways. The present work did not include such pathways because although CoCoMac does contain tracer data regarding the presence/absence of thalamic connections, it does not provide connection weights, which are critical to constraining the dynamics of the model. We have validated the tractography methodologies used here for estimating cortical connection strengths against available tracer data (Shen et al., 2019); however, validated methods for tractography in subcortical regions in macaques do not currently exist. Nonetheless, a lack of subcortical considerations does not diminish the evidence of the rapid communication between the hippocampal and oculomotor systems via cortical routes, within the time window of a typical gaze fixation.

There may be a number of different models that could have adequately addressed our questions of interest here. Testing the performance of multiple models was beyond the scope of the current investigation; rather, our goal was to provide insights to guide theory and future empirical studies regarding how neural responses from the HC and MTL may influence neural responses within regions of the oculomotor system. However, it remains unclear which of our model parameters were necessary for our observations of rapid signal propagation between the systems, limiting the biological interpretation regarding relevant model parameters. In particular, although our model of choice exhibits some of the emergent properties of spontaneous activity (i.e., resting-state networks; Spiegler et al 2016), it remains to be seen how the presence of spontaneous activity may affect signal propagation between the memory and oculomotor systems. Future work examining the effects of model parameters (e.g., adding local noise to generate spontaneous activity) or model type (e.g., comparison to diffusion models) is still needed.

Neuropsychological, neuroimaging, and neurophysiological studies provide important information regarding the representational content that is supported by distinct regions of the brain. A network analysis approach can be instrumental in revealing the broad dynamics by which such representational content governs behavior (Mišić, Goñi, Betzel, Sporns, & McIntosh, 2014; Vlachos, Aertsen, & Kumar, 2012). Memory for objects and their spatial relations provide rapid guidance for gaze prioritization and accurate targeting for foveation. Disruptions to the HC/MTL result in an altered rate and pattern of visual exploration (Hannula et al., 2007; Olsen et al., 2015), consistent with the dynamics of our lesion models. The contribution of HC/MTL is not considered in most models of oculomotor guidance and control (Hamker, 2006; Itti & Koch, 2000) (Belopolsky, 2015). The present work therefore calls for a reconsideration of the neural architecture that supports oculomotor guidance: the HC/MTL provides information to guide visual exploration across space and time. Exciting empirical research shows that functional activity and neural oscillations in the HC/MTL are modified through gaze behavior (Hoffman et al., 2013; Killian et al., 2015; Leonard et al., 2015; Z.-X. Liu et al., 2017). Empirical research is now needed to explore the predictions made by the model here; namely, that the information and signal emanating from the HC/MTL directly influences activity within the oculomotor system and can determine the targets of foveation.

## Methods

Large-scale network dynamics were simulated using TheVirtualBrain (TVB; thevirtualbrain.org) software platform. The connectome-based model represented each node of the network as a neural mass, all coupled together according to a structural connectivity matrix which constrains the spatial and temporal interactions of the system (Breakspear, 2017; Gustavo Deco, Jirsa, & McIntosh, 2011).

### Data

A macaque network with 77 nodes of a single hemisphere was defined using the FV91 parcellation (Felleman & Van Essen, 1991) and its structural connectivity was queried using the CoCoMac database of tract tracing studies (cocomac.g-node.org) (Bakker, Wachtler, & Diesmann, 2012; Stephan et al., 2001). A review of the extant literature was also performed to ensure the accuracy of anatomical pathways within and across MTL and oculomotor systems (Shen et al., 2016). Self-connections were not included in the connectivity matrix.

As CoCoMac only provides categorical weights for connections (i.e., weak, moderate or strong), we ran probabilistic tractography on diffusion-weighted MR imaging data from 10 male adult macaque monkeys (9 *Macaca mulatta*, 1 *Macaca fascicularis*, age 5.8 ± 1.9 years) using the FV91 parcellation to estimate the fibre tract capacities and tract lengths between regions. Image acquisition, preprocessing and tractography procedures for this particular dataset have been previously described (Shen et al., in press., 2019). Fiber tract capacity estimates (i.e., ‘weights’) between each ROI pair were computed as the number of streamlines detected between them, normalized by the total number of streamlines that were seeded. Connectivity weight estimates were averaged across animals and applied to the tracer network, keeping only the connections that appear in the tracer network. The resulting structural connectome was therefore directed, as defined by the tracer data, and fully weighted, as estimated from tractography. Tract lengths were also estimated using probabilistic tractography.

### Node dynamics

The dynamics of each node in the macaque network were given by the following generic 2-dimensional planar oscillator equations:

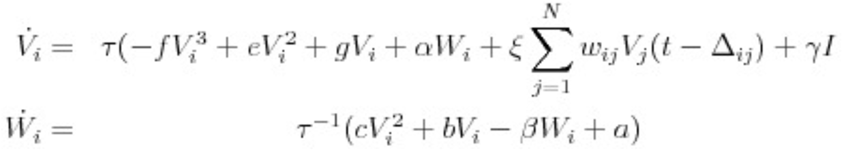

where the fast variable *V* represents mean subthreshold neural activity (i.e., local field potential) at node *i*, *W* is a slower timescale recovery variable; the differential time constants of *V* vs. *W* are controlled by the time scale separation parameter τ.

The local coupling is scaled by *g*, while the global connectivity scaling factor ξ acts on all incoming connections to each node, which are also weighted individually by the connectivity weights matrix *w* (as described above). Exogenous stimulation currents of interest in the present study enter the system through the input variable *I*. Transmission between network nodes was constrained according to the conduction delays matrix *Δ* = *L/v*, where *L* is a matrix of inter-regional tract lengths and *v* is axonal conduction velocity. As in Spiegler et al. (Spiegler et al., 2016), cubic, quadratic, and linear coefficients for *V* and *W* were set such that the dynamics reduce to a classic Fitzhugh-Nagumo system. Additional model parameters are listed in Supplementary Table 1.

Brain dynamics operate near criticality (Ghosh et al., 2008). In this state, the nodes will naturally oscillate with constant magnitude. Setting the local parameter *g* so that the system operates near criticality will allow the node to respond with a strong amplitude, and a longer lasting oscillation. If far from the critical point, the amplitude responses will be weak, slow, and fade quickly, and if spreading within a network, the excitation will decay quickly as it travels. Given our network’s structure, re-entry points allow a node to be re-stimulated, making the excitation last longer and travel farther through the network (Spiegler et al., 2016). Following Spiegler et al. (Spiegler et al., 2016), we set the model parameter *g* was to −0.1 such that the system operated close to the criticality by dampening local excitability. The system of delay-differential equations shown above were solved numerically using a Heun Deterministic integration scheme, with step size dt=0.1 ms.

### Model tuning & stimulation parameters

Simulations were run for 7000 ms, with stimulus onset occurring after 5000 ms to allow for settling of the initial transient resulting from randomly specified initial conditions. A single pulsed stimulus was used, with duration of 100 ms. To determine when nodes became active following stimulation, we first computed the envelope of each node’s timeseries using a Hilbert transform. Each node’s baseline activity was taken as the mean amplitude of the envelope in the 200 ms prior to stimulation. The activation threshold of each node was defined as the baseline activity ± 2 std and activation time of each node was taken as the time its envelope amplitude surpassed the activation threshold.

To create a biologically realistic model, we stimulated V1 to find activation times of the following areas: V1, V2, V3, V4, middle temporal and medial superior temporal. Conduction velocity (*v*) was set to 3.0 m/s, and was within the range of conduction velocities estimated in empirical studies of the macaque brain (Caminiti et al., 2013; Girard, Hupé, & Bullier, 2001). Activation times following V1 stimulation were compared to available empirical data (Schmolesky et al., 2017) and relevant model parameters were adjusted accordingly. Global coupling *ξ* was set to 0.012 and stimulus weighting (*γ*) was set to 0.03 so that simulated response times of visual areas following V1 stimulation exhibited a pattern of activations resembling the known hierarchical processing organization of the visual system (V2: 4 ms, V3: 4 ms, V4: 8 ms, MT: 9 ms, MSTl: 37 ms, MSTd: 47 ms, FEF: 99 ms). Differences with empirical activation times (e.g., MST and FEF) may be due 1) a lack of subcortical-cortical pathways in our model; and 2) the use of the same conduction velocity for all connections.

The same model and stimulation parameters were then used to stimulate the subregions of the hippocampus (CA3, CA1, subiculum, pre-subiculum, para-subiculum), enthorinal cortex (ERC), areas 35 and 36 of the perirhinal cortex (PRC), and areas TF and TH of the parahippocampal cortex (PHC), to look for the activations of nodes whose pathways may serve to mediate the exchange of information between the memory and oculomotor systems (Shen et al., 2016). These nodes of interest included areas V2, V3, V4, VP, MT, MSTd, PO, MIP, FST, 7a, granular insular cortex, anterior cingulate cortex, 46, 12, proisocortex, 5, 10, 11, orbitofrontal area 13, orbitofrontal area 14, 23, 25, 32,and 11. We further examined whether activation was observed in regions important for oculomotor guidance, including the lateral intraparietal area (area LIP), the dorsolateral prefrontal cortex (area 46), anterior cingulate cortex (area 24), and the frontal eye fields (FEF).

### Lesion models

Lesions of particular HC and MTL subregions were simulated by removing their afferent and efferent connections to the rest of the network. Stimulations of other HC and MTL sites were repeated on these lesion models.

### Code availability

Simulations were carried out using the command-line version of TheVirtualBrain (TVB) software package in Python, which is available for download at http://thevirtualbrain.org. The customized TVB code for the simulations presented here is available upon request.

## Acknowledgements

This work was supported by funding awarded to J.D.R. from the Natural Sciences and Engineering Research Council of Canada (NSERC; 482639).

