## Supplementary Materials for "Modelling the influence of the hippocampal memory system on the oculomotor system"

**Supplementary Table 1. Additional Generic 2D oscillator model parameters**

| **Parameter** | **Value** | **Description** |
|---|---|---|
| $d$ | 0.07674 | Temporal scale factor |
| $\tau$ | 1 | Time-scale hierarchy parameter |
| $f$ | 1 | Coefficient for fast variable cubic self-feedback term |
| $e$ | 0 | Coefficient for fast variable quadratic self-feedback term |
| $g$ | -0.1 | Coefficient of fast variable linear self-feedback term |
| $\alpha$ | 1 | Coefficient for linear input term from slow to fast variable |
| $\gamma$ | 1 | Additional scaling parameter for fast variable constant input *I* and long-range inputs |
| $c$ | 0 | Coefficient for quadratic input term from fast variable to slow variable |
| $b$ | -12.3038 | Coefficient for linear input term from fast variable to slow variable |
| $\beta$ | 0 | Coefficient for slow variable linear self-feedback term |
| $a$ | 0 | Slow variable constant input term |
| $I$ | 0 | Fast variable constant input term |

**Supplementary Table 2. Activation times (ms) following simulated stimulation of hippocampal subfields and medial temporal lobe regions for each node in the model.** Observation nodes are presented in alphanumeric order. Observations nodes of interest noted in the main manuscript (HC/MTL regions, oculomotor nodes, and regions that are involved in the shortest paths between HC/MTL and oculomotor nodes) are bolded. S= subiculum, PrS = pre-subiculum, PaS=para-subiculum; ERC = entorhinal cortex; 35/36 = perirhinal cortex; TF/TH = parahippocampal cortex. *0* = stimulation onset; N/A = no response observed.

| | | **Stimulated Node** | | | | | | | | | |
|---|---|---|---|---|---|---|---|---|---|---|---|
| | | **CA3** | **CA1** | **S** | **PrS** | **PaS** | **ERC** | **35** | **36** | **TF** | **TH** |
| **Observation Node** | 1 | N/A | 676 | N/A | 157 | N/A | 221 | 125 | 145 | 312 | 174 |
| | 2 | N/A | 645 | N/A | 119 | N/A | 372 | 151 | 206 | 251 | 153 |
| | 4 | N/A | 534 | N/A | 141 | N/A | 265 | 103 | 115 | 262 | 152 |
| | **5** | N/A | 322 | 50 | 92 | 555 | 249 | 129 | 141 | 69 | 64 |
| | 6 | N/A | 239 | 630 | 25 | 816 | 115 | 22 | 15 | 82 | 39 |
| | 9 | N/A | 109 | 192 | 14 | 173 | 15 | 12 | 15 | 18 | 17 |
| | **10** | N/A | 59 | 24 | 83 | 106 | 12 | 10 | 12 | 81 | 20 |
| | **11** | N/A | 12 | 68 | 74 | 166 | 10 | 7 | 8 | 38 | 48 |
| | **12** | N/A | 96 | 186 | 93 | 285 | 15 | 9 | 10 | 24 | 34 |
| | **13** | N/A | 10 | 18 | 62 | 132 | 8 | 7 | 11 | 19 | 19 |
| | **14** | N/A | 9 | 27 | 57 | 76 | 9 | 9 | 11 | 13 | 20 |
| | **23** | N/A | 165 | 384 | 12 | 231 | 43 | 37 | 52 | 24 | 13 |
| | **24** | N/A | 107 | 250 | 55 | 238 | 24 | 22 | 25 | 22 | 16 |
| | **25** | 435 | 6 | 19 | 35 | 57 | 6 | 11 | 8 | 9 | 12 |
| | 29 | 98 | 20 | 133 | 32 | 42 | 10 | 21 | 19 | 0 | 0 |
| | 30 | 336 | 55 | 265 | 61 | 118 | 14 | 127 | 128 | 0 | 2 |
| | **32** | N/A | 38 | 63 | 88 | 89 | 10 | 13 | 14 | 14 | 17 |
| | **35** | 567 | 6 | 21 | 49 | 50 | 0 | *0* | 0 | 9 | 19 |
| | **36** | 317 | 1 | 14 | 10 | 6 | 0 | 0 | *0* | 6 | 14 |
| | 45 | N/A | 172 | 334 | 84 | 464 | 63 | 14 | 15 | 23 | 88 |
| | **46** | N/A | 84 | 135 | 15 | 187 | 12 | 9 | 11 | 17 | 23 |
| | 3a | N/A | 631 | N/A | 143 | N/A | 409 | 89 | 124 | 292 | 190 |
| | 3b | N/A | 767 | N/A | 333 | N/A | 184 | 113 | 126 | 482 | 236 |
| | **7a** | N/A | 381 | 202 | 336 | 795 | 256 | 53 | 52 | 103 | 59 |
| | 7b | N/A | 385 | 566 | 152 | N/A | 318 | 101 | 124 | 155 | 132 |
| | A1 | N/A | N/A | N/A | N/A | N/A | 48 | 18 | 24 | N/A | 64 |
| | AII | N/A | N/A | N/A | N/A | N/A | 108 | 20 | 18 | 409 | 165 |
| | AITd | N/A | 159 | 437 | 210 | 710 | 469 | 409 | 341 | 35 | 200 |
| | AITv | 365 | 23 | 129 | 40 | 146 | 128 | 134 | 83 | 0 | 11 |
| | **CA1** | 137 | *0* | 0 | 0 | 46 | 0 | 8 | 2 | 0 | 0 |
| | **CA3** | *0* | 48 | 31 | 37 | 27 | 0 | 21 | 20 | 48 | 1 |
| | CITd | N/A | 226 | 639 | 270 | 604 | 604 | 381 | 370 | 56 | 121 |
| | CITv | N/A | 34 | 179 | 57 | 327 | 160 | 134 | 103 | 0 | 59 |
| | CM | N/A | N/A | N/A | N/A | N/A | 136 | 40 | 40 | 805 | 83 |

| | | | | | | | | | | | |
|---|---|---|---|---|---|---|---|---|---|---|---|
| | DP | N/A | 362 | N/A | 355 | 678 | 368 | 172 | 173 | 95 | 105 |
| | **ERC** | 169 | 40 | 0 | 4 | 0 | *0* | 0 | 0 | 7 | 13 |
| | **FEF** | N/A | 217 | 452 | 68 | 500 | 79 | 19 | 15 | 34 | 71 |
| | **FST** | 186 | 47 | 147 | 52 | 79 | 6 | 1 | 3 | 0 | 6 |
| | G | N/A | 470 | N/A | 150 | N/A | 278 | 66 | 69 | 366 | 214 |
| | Id | N/A | 90 | 241 | 120 | 268 | 12 | 7 | 7 | 32 | 19 |
| | **Ig** | N/A | 11 | 63 | 75 | 189 | 11 | 4 | 6 | 21 | 13 |
| | L | N/A | N/A | N/A | N/A | N/A | 104 | 21 | 17 | 459 | 293 |
| | **LIP** | N/A | 535 | N/A | 445 | N/A | 561 | 233 | 230 | 176 | 144 |
| | MDP | N/A | 548 | 353 | 307 | 874 | 544 | 263 | 276 | 211 | 178 |
| | **MIP** | N/A | 237 | 523 | 215 | 296 | 405 | 113 | 126 | 70 | 48 |
| | **MSTd** | N/A | 286 | 664 | 287 | 494 | 33 | 23 | 23 | 94 | 53 |
| | MSTl | N/A | 145 | 329 | 151 | 238 | 19 | 11 | 12 | 49 | 29 |
| | **MT** | N/A | 260 | 857 | 272 | 483 | 189 | 107 | 23 | 36 | 83 |
| | Pa | N/A | N/A | N/A | N/A | N/A | 33 | 13 | 18 | 525 | 130 |
| | PAC | 60 | 43 | 41 | 42 | 37 | 0 | 0 | 0 | 49 | 4 |
| | **PaS** | 350 | 17 | 96 | 28 | *0* | 50 | 15 | 0 | 0 | 32 |
| | PIP | N/A | 250 | 743 | 228 | 315 | 454 | 134 | 147 | 76 | 52 |
| | Pir | 166 | 64 | 23 | 37 | 24 | 0 | 24 | 24 | 62 | 11 |
| | PITd | N/A | 217 | 637 | 226 | 360 | 460 | 153 | 171 | 52 | 64 |
| | PITv | N/A | 177 | 507 | 224 | 538 | 485 | 339 | 315 | 37 | 114 |
| | **PO** | 656 | 208 | 593 | 190 | 248 | 355 | 99 | 107 | 60 | 39 |
| | **Proisocortex** | N/A | 52 | 96 | 12 | 48 | 3 | 1 | 4 | 29 | 20 |
| | Prostriata | N/A | N/A | N/A | N/A | N/A | N/A | N/A | N/A | N/A | N/A |
| | **PrS** | 84 | 23 | 119 | *0* | 25 | 6 | 13 | 11 | 0 | 0 |
| | Ri | N/A | 534 | N/A | 702 | N/A | 208 | 68 | 82 | 351 | 249 |
| | RL | N/A | 267 | N/A | N/A | N/A | 38 | 9 | 8 | 457 | 34 |
| | **S** | 115 | 17 | *0* | 26 | 37 | 0 | 16 | 10 | 0 | 0 |
| | SII | N/A | 124 | 397 | 140 | N/A | 74 | 7 | 9 | 46 | 123 |
| | SMA | N/A | 377 | 641 | 153 | 957 | 174 | 139 | 118 | 172 | 98 |
| | STPa | N/A | 46 | 110 | 59 | 104 | 4 | 0 | 0 | 2 | 9 |
| | STPp | N/A | 252 | N/A | 369 | N/A | 89 | 10 | 8 | 75 | 74 |
| | **TF** | 137 | 0 | 20 | 0 | 44 | 11 | 12 | 6 | *0* | 0 |
| | **TH** | 7 | 0 | 29 | 0 | 0 | 20 | 29 | 18 | 0 | *0* |
| | V1 | N/A | 217 | 670 | 197 | 261 | 399 | 117 | 125 | 64 | 40 |
| | **V2** | 215 | 56 | 274 | 48 | 66 | 139 | 16 | 17 | 6 | 0 |
| | **V3** | N/A | 235 | 733 | 224 | 313 | 389 | 126 | 137 | 60 | 52 |
| | V3A | N/A | 260 | 841 | 257 | 409 | 398 | 154 | 163 | 68 | 68 |
| | **V4** | N/A | 76 | 353 | 103 | 244 | 280 | 147 | 142 | 5 | 15 |
| | V4t | N/A | 273 | 936 | 280 | 495 | 369 | 175 | 149 | 66 | 88 |
| | VIP | N/A | 506 | N/A | 391 | N/A | 508 | 195 | 202 | 160 | 116 |
| | VOT | N/A | 231 | 665 | 245 | 401 | 489 | 177 | 194 | 57 | 75 |
| | **VP** | 753 | 55 | 266 | 86 | 259 | 223 | 98 | 101 | 2 | 41 |

**Supplementary Table 3. Changes in activation times (ms) for nodes of interest following a lesion of CA1 and simulated stimulation of hippocampal subfields and medial temporal lobe regions.** Values were determined by subtracting the intact stimulation times (Table 1) from lesioned activation times, such that slower responses are positive while faster responses are negative.

| | | Stimulated Node | | | | | | | | | |
|---|---|---|---|---|---|---|---|---|---|---|---|
| | | **CA3** | **CA1** | **S** | **PrS** | **PaS** | **ERC** | **35** | **36** | **TF** | **TH** |
| **Observation Node** | **CA3** | *0* | | -4 | -2 | 0 | 0 | 0 | 0 | -5 | -1 |
| | **CA1** | | | | | | | | | | |
| | **S** | -10 | | *0* | -7 | -4 | 0 | -1 | -1 | 0 | 0 |
| | **PrS** | -4 | | -3 | *0* | -1 | -2 | -1 | -1 | 0 | 0 |
| | **PaS** | -32 | | 106 | -7 | *0* | -8 | -1 | 0 | 0 | -8 |
| | **ERC** | -3 | | 0 | -1 | 0 | *0* | 0 | 0 | -1 | -1 |
| | **35** | N/A | | -1 | 0 | 0 | 0 | *0* | 0 | -1 | -3 |
| | **36** | 55 | | -2 | -1 | 0 | 0 | 0 | *0* | -1 | -2 |
| | **TF** | -19 | | 214 | 0 | -9 | -3 | -2 | -1 | *0* | 0 |
| | **TH** | 0 | | 327 | 0 | 0 | -3 | -4 | -2 | 0 | *0* |
| | **5** | N/A | | 1 | 0 | 2 | 0 | 0 | -1 | 0 | 0 |
| | **10** | N/A | | -1 | -6 | 1 | 0 | 0 | 0 | -7 | -1 |
| | **11** | N/A | | 25 | 11 | 3 | 0 | 0 | 0 | 1 | -3 |
| | **12** | N/A | | 10 | -6 | 3 | 0 | 0 | 1 | -2 | -2 |
| | **13** | N/A | | 1 | 17 | 2 | 0 | 0 | 0 | 0 | -1 |
| | **14** | N/A | | 0 | 17 | 1 | 0 | 0 | 0 | 0 | -1 |
| | **23** | N/A | | N/A | 0 | 0 | 0 | 0 | 0 | 0 | 0 |
| | **25** | 63 | | -2 | 14 | 2 | 0 | 0 | 0 | 0 | -1 |
| | **32** | N/A | | 0 | -4 | 1 | 0 | 0 | 0 | 0 | 0 |
| | **Ig** | N/A | | 22 | 28 | 4 | 0 | 0 | 0 | 1 | 0 |
| | **Pro** | N/A | | 3 | 0 | 1 | 1 | 0 | 0 | 0 | 0 |
| | **7a** | N/A | | -2 | 1 | -1 | -2 | -1 | -1 | -1 | 0 |
| | **FST** | -5 | | 13 | -3 | -1 | 0 | 0 | 0 | 0 | 0 |
| | **MIP** | N/A | | N/A | -4 | -1 | 4 | -1 | -1 | -2 | -1 |
| | **PO** | -11 | | N/A | -6 | -2 | 3 | -2 | -1 | -2 | -1 |
| | **MSTd** | N/A | | N/A | 1 | 2 | 0 | 0 | 0 | -1 | 0 |
| | **MT** | N/A | | N/A | 0 | -2 | -1 | -1 | -1 | -1 | 0 |
| | **V4** | N/A | | N/A | -4 | -2 | 11 | -2 | -2 | -1 | -1 |
| | **VP** | -19 | | N/A | -5 | -3 | 8 | -2 | -2 | 0 | -2 |
| | **V3** | N/A | | N/A | -5 | -2 | 4 | -1 | -3 | -2 | 0 |
| | **V2** | -3 | | N/A | -4 | -1 | -1 | 0 | 0 | -1 | 0 |
| | **24** | N/A | | 33 | -1 | 3 | 0 | 0 | 0 | 0 | 0 |
| | **46** | N/A | | 1 | 0 | 2 | 0 | 0 | 0 | -1 | -1 |
| | **FEF** | N/A | | 84 | -3 | 5 | 0 | 0 | 0 | -1 | -1 |
| | **LIP** | N/A | | N/A | 0 | N/A | 0 | -2 | -2 | -3 | -2 |

**Supplementary Table 4. Changes in activation times (ms) for nodes of interest following a lesion of PrS and simulated stimulation of hippocampal subfields and medial temporal lobe regions.** Conventions as in Supplementary Table 3.

| | | Stimulated Node | | | | | | | | | |
|---|---|---|---|---|---|---|---|---|---|---|---|
| | | **CA3** | **CA1** | **S** | **PrS** | **PaS** | **ERC** | **35** | **36** | **TF** | **TH** |
| **Observation Node** | **CA3** | *0* | -3 | 0 | | -1 | 0 | 0 | 0 | -6 | -1 |
| | **CA1** | -17 | *0* | 0 | | -9 | 0 | -1 | -2 | 0 | 0 |
| | **S** | -9 | 0 | *0* | | -3 | 0 | -1 | 0 | 0 | 0 |
| | **PrS** | | | | | | | | | | |
| | **PaS** | -20 | 1 | -3 | | *0* | -4 | 0 | 0 | 0 | -8 |
| | **ERC** | 38 | -3 | 0 | | 0 | *0* | 0 | 0 | -1 | -3 |
| | **35** | N/A | 0 | 0 | | 0 | 0 | *0* | 0 | 0 | -3 |
| | **36** | 38 | 0 | 0 | | 0 | 0 | 0 | *0* | -1 | -2 |
| | **TF** | -14 | 0 | 1 | | -7 | -2 | -1 | 0 | *0* | 0 |
| | **TH** | -1 | 0 | -4 | | 0 | -4 | -5 | -2 | 0 | *0* |
| | **5** | N/A | 3 | 1 | | 12 | 5 | 0 | -1 | 0 | 0 |
| | **10** | N/A | -1 | 0 | | 0 | 0 | 0 | 0 | -5 | -1 |
| | **11** | N/A | 0 | 0 | | 2 | 0 | 0 | 0 | -2 | -5 |
| | **12** | N/A | 0 | 1 | | 2 | 0 | 0 | 1 | -1 | -3 |
| | **13** | N/A | 0 | 0 | | 1 | 0 | 0 | 0 | -1 | -1 |
| | **14** | N/A | 0 | 0 | | 1 | 0 | 0 | 0 | 0 | -1 |
| | **23** | N/A | 4 | 1 | | 5 | 0 | -1 | 0 | 0 | 0 |
| | **25** | 8 | 1 | 0 | | 1 | 0 | 0 | 0 | 1 | -1 |
| | **32** | N/A | 0 | 0 | | 0 | 0 | 0 | 0 | 0 | 0 |
| | **Ig** | N/A | 0 | 1 | | 2 | 0 | 0 | 0 | 1 | 0 |
| | **Pro** | N/A | 0 | 1 | | 1 | 1 | 0 | 0 | 0 | 0 |
| | **7a** | N/A | -2 | 0 | | 1 | -2 | -1 | -1 | -1 | -1 |
| | **FST** | -6 | -1 | -1 | | -1 | 0 | 0 | 0 | 0 | 0 |
| | **MIP** | N/A | -2 | -5 | | -2 | 3 | -2 | -1 | -1 | 2 |
| | **PO** | -13 | -2 | -6 | | -3 | 2 | -2 | -1 | -1 | -2 |
| | **MSTd** | N/A | 0 | 4 | | 2 | 0 | 0 | 0 | 0 | -1 |
| | **MT** | N/A | -1 | -9 | | -3 | -1 | -1 | -1 | 0 | -1 |
| | **V4** | N/A | -1 | -3 | | -3 | 0 | -2 | -2 | 0 | -1 |
| | **VP** | -16 | -2 | -3 | | -3 | 0 | -2 | -2 | 0 | -3 |
| | **V3** | N/A | -2 | -8 | | -3 | 0 | -1 | -3 | -1 | -1 |
| | **V2** | -4 | -1 | -4 | | -2 | -1 | 0 | -1 | -1 | 0 |
| | **24** | N/A | 1 | 3 | | 3 | 1 | 0 | 0 | 0 | -1 |
| | **46** | N/A | -1 | 1 | | 5 | 0 | 0 | 0 | -1 | -1 |
| | **FEF** | N/A | 0 | 4 | | 18 | 0 | 0 | 0 | 0 | -4 |
| | **LIP** | N/A | -4 | N/A | | N/A | -5 | -2 | -2 | -3 | -3 |

**Supplementary Table 5. Changes in activation times (ms) for nodes of interest following a lesion of all hippocampal subfields and simulated stimulation of medial temporal lobe regions.** Conventions as in Supplementary Table 3.

| | | Stimulated Node | | | | | | | | | |
|---|---|---|---|---|---|---|---|---|---|---|---|
| | | **CA3** | **CA1** | **S** | **PrS** | **PaS** | **ERC** | **35** | **36** | **TF** | **TH** |
| **Observation Node** | **CA3** | | | | | | | | | | |
| | **CA1** | | | | | | | | | | |
| | **S** | | | | | | | | | | |
| | **PrS** | | | | | | | | | | |
| | **PaS** | | | | | | | | | | |
| | **ERC** | | | | | | *0* | 0 | 0 | -3 | -4 |
| | **35** | | | | | | 0 | *0* | 0 | -1 | -5 |
| | **36** | | | | | | 0 | 0 | *0* | -2 | -3 |
| | **TF** | | | | | | -4 | -3 | -4 | *0* | 0 |
| | **TH** | | | | | | -6 | -7 | -4 | 0 | *0* |
| | **5** | | | | | | 14 | -1 | -2 | 0 | 0 |
| | **10** | | | | | | 0 | 0 | 0 | -10 | -2 |
| | **11** | | | | | | 0 | 0 | 0 | 0 | -7 |
| | **12** | | | | | | 0 | 0 | 1 | -3 | -5 |
| | **13** | | | | | | 0 | 0 | 0 | -1 | -2 |
| | **14** | | | | | | 0 | 0 | 0 | 0 | -2 |
| | **23** | | | | | | 0 | -1 | 0 | 0 | -1 |
| | **25** | | | | | | 0 | 0 | 0 | 0 | -1 |
| | **32** | | | | | | 0 | 0 | 0 | -1 | -1 |
| | **Ig** | | | | | | 0 | 0 | 0 | 1 | 0 |
| | **Pro** | | | | | | 1 | 0 | 0 | 0 | -1 |
| | **7a** | | | | | | 0 | -1 | -1 | -2 | -1 |
| | **FST** | | | | | | 0 | 1 | 0 | 0 | -1 |
| | **MIP** | | | | | | 12 | -3 | -2 | -2 | -2 |
| | **PO** | | | | | | 7 | -3 | -2 | -3 | -2 |
| | **MSTd** | | | | | | 0 | 0 | 0 | -2 | -1 |
| | **MT** | | | | | | -2 | -2 | -1 | -1 | -1 |
| | **V4** | | | | | | 14 | -3 | -3 | -1 | -1 |
| | **VP** | | | | | | 8 | -4 | -3 | -1 | -3 |
| | **V3** | | | | | | 7 | -3 | -5 | -3 | -2 |
| | **V2** | | | | | | -1 | 0 | -1 | -1 | 0 |
| | **24** | | | | | | 1 | 0 | 0 | 0 | -1 |
| | **46** | | | | | | 0 | 0 | 0 | -1 | -2 |
| | **FEF** | | | | | | 0 | 0 | 0 | -2 | -5 |
| | **LIP** | | | | | | 1 | -4 | -4 | -4 | -4 |

**Supplementary Table 6. Changes in activation times (ms) for nodes of interest following a lesion of the ERC and simulated stimulation of hippocampal subfields and medial temporal lobe regions.** Conventions as in Supplementary Table 3.

| | | Stimulated Node | | | | | | | | | |
|---|---|---|---|---|---|---|---|---|---|---|---|
| | | CA3 | CA1 | S | PrS | PaS | ERC | 35 | 36 | TF | TH |
| Observation Node | CA3 | *0* | -8 | 127 | -2 | 6 | | 54 | 51 | -6 | -1 |
| | CA1 | -1 | *0* | 0 | 0 | 0 | | -1 | -2 | 0 | 0 |
| | S | -3 | 0 | *0* | -2 | 2 | | -2 | -1 | 0 | 0 |
| | PrS | 0 | 0 | 14 | *0* | -1 | | -1 | -1 | 0 | 0 |
| | PaS | -3 | 0 | 0 | 0 | *0* | | -1 | 0 | 0 | 0 |
| | ERC | | | | | | | | | | |
| | 35 | -22 | 0 | -1 | 4 | 34 | | *0* | 0 | -1 | -3 |
| | 36 | -17 | -1 | -2 | -1 | 0 | | 0 | *0* | -1 | -1 |
| | TF | -1 | 0 | 1 | 0 | -1 | | -1 | 0 | *0* | 0 |
| | TH | 0 | 0 | 0 | 0 | 0 | | -2 | -1 | 0 | *0* |
| | 5 | N/A | 0 | 0 | 0 | 1 | | -2 | -4 | 0 | 0 |
| | 10 | N/A | -1 | -1 | 4 | 75 | | -1 | -1 | -2 | 0 |
| | 11 | N/A | 0 | 15 | 2 | 139 | | -1 | -1 | -1 | -2 |
| | 12 | N/A | -1 | 107 | -2 | 267 | | 0 | 0 | -1 | -1 |
| | 13 | N/A | 0 | 1 | 3 | 108 | | 0 | -1 | 0 | -1 |
| | 14 | N/A | 0 | 1 | 3 | 104 | | -1 | -1 | 0 | -1 |
| | 23 | N/A | 0 | -5 | 0 | 1 | | -1 | -1 | 0 | 0 |
| | 25 | 26 | 0 | -3 | 0 | 45 | | -1 | -1 | 0 | -1 |
| | 32 | N/A | 0 | 70 | 20 | 86 | | -1 | -1 | 0 | 0 |
| | Ig | N/A | 0 | 2 | 2 | 28 | | 0 | -1 | 0 | 0 |
| | Pro | N/A | 0 | 98 | 0 | 1 | | -1 | -1 | 0 | 0 |
| | 7a | N/A | -1 | -2 | 0 | 22 | | -1 | 0 | 0 | 0 |
| | FST | -1 | -1 | 78 | -1 | 3 | | -1 | -1 | 0 | 0 |
| | MIP | N/A | 0 | -6 | 0 | 0 | | -1 | 0 | 0 | 0 |
| | PO | 0 | 0 | -10 | 0 | 0 | | -1 | 0 | 0 | 0 |
| | MSTd | N/A | 0 | N/A | 8 | 46 | | 0 | 0 | 0 | 0 |
| | MT | N/A | 0 | 72 | -1 | 7 | | 0 | 0 | 0 | 0 |
| | V4 | N/A | 0 | -3 | 0 | 0 | | 0 | 0 | 0 | 0 |
| | VP | -1 | 0 | -2 | 0 | 0 | | -1 | -1 | 0 | 0 |
| | V3 | N/A | 0 | -17 | 0 | 0 | | 0 | -1 | 0 | 0 |
| | V2 | 0 | 0 | -4 | 0 | 0 | | 0 | 0 | 0 | 0 |
| | 24 | N/A | -1 | 96 | -1 | 29 | | 0 | -1 | 0 | -1 |
| | 46 | N/A | -2 | 101 | 0 | 64 | | 0 | 0 | 0 | 0 |
| | FEF | N/A | -3 | 342 | -2 | 106 | | -1 | 0 | 0 | -1 |
| | LIP | N/A | -1 | N/A | -1 | N/A | | -1 | -1 | -1 | 0 |

**Supplementary Table 7. Changes in activation times (ms) for nodes of interest following a combined lesion of areas TH and TF and simulated stimulation of hippocampal subfields and medial temporal lobe regions.** Conventions as in Supplementary Table 3.

| | | Stimulated Node | | | | | | | | | |
|---|---|---|---|---|---|---|---|---|---|---|---|
| | | CA3 | CA1 | S | PrS | PaS | ERC | 35 | 36 | TF | TH |
| Observation Node | CA3 | *0* | 115 | -3 | -1 | -4 | 0 | 0 | 1 | | |
| | CA1 | N/A | *0* | 0 | 0 | 45 | 0 | -1 | -2 | | |
| | S | N/A | 120 | *0* | 17 | -5 | 0 | -4 | -2 | | |
| | PrS | N/A | 185 | -29 | *0* | 101 | -6 | -3 | -2 | | |
| | PaS | N/A | 23 | 16 | 33 | *0* | -16 | -4 | 0 | | |
| | ERC | N/A | -7 | 0 | -3 | 0 | *0* | 0 | 0 | | |
| | 35 | N/A | 0 | 0 | -5 | 1 | 0 | *0* | 0 | | |
| | 36 | N/A | -1 | -1 | -1 | 0 | 0 | 0 | *0* | | |
| | TF | | | | | | | | | | |
| | TH | | | | | | | | | | |
| | 5 | N/A | N/A | 2 | 1 | N/A | 3 | -1 | -3 | | |
| | 10 | N/A | -4 | -1 | -4 | 10 | 0 | 0 | 0 | | |
| | 11 | N/A | 0 | 0 | -7 | 4 | 0 | 0 | 0 | | |
| | 12 | N/A | 1 | 1 | -5 | 35 | 0 | 0 | 1 | | |
| | 13 | N/A | 0 | 1 | -3 | 11 | 0 | 0 | 0 | | |
| | 14 | N/A | 0 | 1 | -5 | 3 | 0 | 0 | 0 | | |
| | 23 | N/A | N/A | N/A | 0 | N/A | -1 | -1 | 0 | | |
| | 25 | N/A | 0 | 0 | -1 | 6 | 0 | 0 | 0 | | |
| | 32 | N/A | -1 | 0 | 11 | 6 | 0 | 0 | 0 | | |
| | Ig | N/A | 1 | 3 | 31 | 142 | 0 | 0 | 0 | | |
| | Pro | N/A | 0 | 1 | 0 | 2 | 1 | 0 | 0 | | |
| | 7a | N/A | N/A | 5 | N/A | N/A | -4 | -2 | -2 | | |
| | FST | N/A | 68 | 12 | 96 | 130 | 0 | 2 | 1 | | |
| | MIP | N/A | N/A | N/A | 349 | N/A | 27 | -8 | -8 | | |
| | PO | N/A | N/A | N/A | N/A | N/A | 19 | -9 | -7 | | |
| | MSTd | N/A | N/A | N/A | N/A | N/A | 1 | 0 | 0 | | |
| | MT | N/A | N/A | N/A | N/A | N/A | -4 | -4 | -1 | | |
| | V4 | N/A | N/A | N/A | N/A | N/A | 67 | -7 | 0 | | |
| | VP | N/A | N/A | N/A | N/A | N/A | -4 | -4 | -1 | | |
| | V3 | N/A | N/A | N/A | N/A | N/A | 17 | -10 | -11 | | |
| | V2 | N/A | N/A | N/A | N/A | N/A | 3 | -1 | -1 | | |
| | 24 | N/A | 46 | 21 | 3 | 412 | 1 | 1 | 0 | | |
| | 46 | N/A | 7 | 1 | -1 | 53 | 0 | 0 | 0 | | |
| | FEF | N/A | 69 | 35 | -5 | N/A | 1 | 0 | 0 | | |
| | LIP | N/A | N/A | N/A | N/A | N/A | 1 | -9 | -8 | | |

**Supplementary Table 8. Changes in activation times (ms) for nodes of interest following a combined lesion of areas 35 and 36 and simulated stimulation of hippocampal subfields and medial temporal lobe regions.** Conventions as in Supplementary Table 3.

| | | Stimulated Node | | | | | | | | | |
|---|---|---|---|---|---|---|---|---|---|---|---|
| | | CA3 | CA1 | S | PrS | PaS | ERC | 35 | 36 | TF | TH |
| Observation Node | CA3 | *0* | -5 | -3 | -1 | -2 | 0 | | | -3 | 0 |
| | CA1 | -2 | *0* | 0 | 0 | -1 | 0 | | | 0 | 0 |
| | S | -3 | 0 | *0* | -1 | -1 | 0 | | | 0 | 0 |
| | PrS | 0 | 0 | -1 | *0* | 0 | -2 | | | 0 | 0 |
| | PaS | -22 | 1 | -7 | 0 | *0* | 34 | | | 0 | 0 |
| | ERC | -19 | 19 | 0 | -3 | 0 | *0* | | | -2 | -2 |
| | 35 | | | | | | | | | | |
| | 36 | | | | | | | | | | |
| | TF | -1 | 0 | 0 | 0 | -1 | -1 | | | *0* | 0 |
| | TH | 0 | 0 | 0 | 0 | 0 | -2 | | | 0 | *0* |
| | 5 | N/A | -1 | 0 | -1 | -5 | 18 | | | 0 | 0 |
| | 10 | N/A | 5 | 0 | -3 | 22 | -2 | | | 5 | -1 |
| | 11 | N/A | 0 | 5 | 1 | 87 | -2 | | | 1 | -3 |
| | 12 | N/A | 17 | 26 | -1 | 158 | -3 | | | -2 | -2 |
| | 13 | N/A | 0 | 1 | -1 | 23 | -1 | | | 0 | -1 |
| | 14 | N/A | 0 | 1 | -3 | 13 | -1 | | | 0 | -1 |
| | 23 | N/A | 0 | -3 | 0 | -1 | -2 | | | 0 | 0 |
| | 25 | 125 | 0 | -1 | -1 | 7 | -1 | | | 1 | -1 |
| | 32 | N/A | 1 | 3 | -4 | 14 | -1 | | | 0 | 0 |
| | Ig | N/A | 0 | -4 | 1 | 45 | -1 | | | 0 | -1 |
| | Pro | N/A | 7 | 8 | 0 | 2 | 0 | | | 1 | 0 |
| | 7a | N/A | 15 | -2 | -1 | 17 | 93 | | | 0 | 0 |
| | FST | -9 | -2 | 2 | -2 | -1 | -1 | | | 0 | 0 |
| | MIP | N/A | 0 | -5 | 0 | 1 | 173 | | | 0 | 0 |
| | PO | 5 | 0 | -10 | 1 | 1 | 158 | | | 0 | 0 |
| | MSTd | N/A | 43 | 285 | 2 | 32 | -1 | | | -1 | -1 |
| | MT | N/A | 3 | 48 | -1 | 10 | 42 | | | 0 | 0 |
| | V4 | N/A | 1 | 2 | 1 | 2 | 19 | | | 0 | 0 |
| | VP | 9 | 0 | 1 | 1 | 1 | 16 | | | 0 | 0 |
| | V3 | N/A | 1 | -15 | 1 | 1 | 142 | | | 0 | 0 |
| | V2 | 1 | 0 | -5 | 0 | 0 | 81 | | | 0 | 0 |
| | 24 | N/A | 1 | 6 | -1 | 6 | -2 | | | 0 | -1 |
| | 46 | N/A | 14 | 11 | 0 | 52 | -1 | | | -1 | -1 |
| | FEF | N/A | 53 | 90 | -2 | 83 | -3 | | | -2 | 0 |
| | LIP | N/A | 4 | N/A | -3 | N/A | N/A | | | -1 | -1 |

**Supplementary Table 9. Changes in activation times (ms) for nodes of interest following a combined lesion of areas V4, 7a, 5 and 23 and simulated stimulation of hippocampal subfields and medial temporal lobe regions.** Conventions as in Supplementary Table 3.

| | Stimulated Node | | | | | | | | | |
|---|---|---|---|---|---|---|---|---|---|---|
| **Observation Node** | **CA3** | **CA1** | **S** | **PrS** | **PaS** | **ERC** | **35** | **36** | **TF** | **TH** |
| **CA3** | *0* | 0 | 0 | 0 | 1 | 0 | 0 | 1 | 1 | 0 |
| **CA1** | -57 | *0* | 0 | 0 | -6 | 1 | -1 | 1 | 0 | 0 |
| **S** | -43 | 0 | *0* | 0 | -2 | 0 | -1 | 0 | 0 | 0 |
| **PrS** | -31 | -1 | -35 | *0* | -2 | -1 | -3 | -2 | 0 | 0 |
| **PaS** | -94 | 0 | -4 | 0 | *0* | -3 | -1 | 0 | 0 | 0 |
| **ERC** | 43 | 2 | 0 | 0 | 0 | *0* | 0 | 0 | 0 | 0 |
| **35** | N/A | 1 | 2 | 2 | 8 | 0 | *0* | 0 | 1 | 0 |
| **36** | 118 | 2 | 1 | 0 | 1 | 0 | 0 | *0* | 0 | 1 |
| **TF** | -54 | 0 | 1 | 0 | -4 | -1 | -2 | 0 | *0* | 0 |
| **TH** | -2 | 0 | -3 | 0 | 0 | -4 | -10 | -4 | 0 | *0* |
| **5** | | | | | | | | | | |
| **10** | N/A | 0 | 0 | -2 | 1 | 0 | 0 | 0 | 0 | 0 |
| **11** | N/A | 0 | 1 | 0 | 10 | 0 | 0 | 0 | 3 | -1 |
| **12** | N/A | 5 | 12 | -1 | 37 | 1 | 0 | 1 | 2 | 1 |
| **13** | N/A | 1 | 1 | -1 | 2 | 0 | 0 | 0 | 0 | 0 |
| **14** | N/A | 1 | 0 | -2 | -2 | 0 | 0 | 0 | 0 | 0 |
| **23** | | | | | | | | | | |
| **25** | -57 | 1 | 1 | 0 | 3 | 0 | 0 | 0 | 1 | 0 |
| **32** | N/A | 0 | -2 | -5 | -4 | 0 | 0 | 0 | 0 | 0 |
| **Ig** | N/A | 0 | -4 | -2 | 11 | 1 | 1 | 0 | 0 | -1 |
| **Pro** | N/A | 2 | 8 | 1 | 8 | 1 | -1 | 0 | 4 | 1 |
| **7a** | | | | | | | | | | |
| **FST** | -12 | 7 | 7 | 1 | 1 | 1 | 4 | 2 | 3 | 1 |
| **MIP** | N/A | -57 | -69 | -39 | -94 | -191 | -55 | -65 | -11 | -4 |
| **PO** | -320 | -36 | -142 | -25 | -54 | -165 | -47 | -51 | -7 | -1 |
| **MSTd** | N/A | -58 | 169 | -58 | -47 | -12 | -6 | -5 | -31 | -19 |
| **MT** | N/A | -39 | -118 | -53 | -102 | -86 | -48 | -9 | -11 | -18 |
| **V4** | | | | | | | | | | |
| **VP** | -432 | -16 | -75 | -22 | -77 | -113 | -53 | -51 | -2 | -8 |
| **V3** | N/A | -49 | -219 | -38 | -81 | -200 | -65 | -72 | -12 | -5 |
| **V2** | -101 | -11 | -59 | -7 | -14 | -68 | -5 | -6 | -1 | 0 |
| **24** | N/A | -12 | -5 | 11 | -24 | -2 | -1 | -2 | -2 | -2 |
| **46** | N/A | 8 | 24 | 1 | 48 | 1 | 1 | 1 | 2 | 1 |
| **FEF** | N/A | 48 | 124 | 7 | 187 | 9 | 2 | 3 | 10 | 6 |
| **LIP** | N/A | -273 | N/A | -197 | N/A | -297 | -109 | -104 | -120 | -79 |
